# Twenty-five years of monitoring reveals that uninterrupted rodent control is the fundamental driver of breeding success in the Galapagos Petrel *Pterodroma phaeopygia*

**DOI:** 10.64898/2026.03.29.715149

**Authors:** Francisco Lopes, James P. Gibbs, Jorge Carrión

## Abstract

The long-standing misconception that the Galapagos petrel (Pterodroma phaeopygia) and the Hawaiian petrel (*Pterodroma sandwichensis*) were conspecific masked the severe vulnerability of the Galápagos population. By the time its distinct status was recognized, the Galápagos petrel was already in marked decline, primarily due to invasive predators. Consequently, sustained rodent control programs have been implemented on Santa Cruz Island. An unintentional one-year failure in rodent control provided a rare quasi-experimental opportunity to quantify the demographic consequences of the invasive black rat predator. During this year, hatching success declined by ∼35% and breeding success by ∼40% relative to long-term means (66% and 62%, respectively), representing a substantial reproductive collapse. Fledging success exhibited a comparatively modest decline (from a long-term mean of 94% to 86% in 2017), suggesting stage-specific vulnerability. These results support the hypothesis that invasive black rats primarily affect early reproductive stages through egg predation and predation on small chicks, while older chicks surpass a critical size threshold that reduces susceptibility. Across the remaining managed years, reproductive metrics exhibited great stability, demonstrating the petrel’s resilience against other environmental or climatic stressors. Our findings provide robust empirical evidence that invasive rodent control is the dominant driver of reproductive success in this endangered seabird. The quasi-experimental failure underscored both the effectiveness and the necessity of continuous predator management, highlighting the severe and immediate consequences of even short-term lapses.

## Introduction

*Pterodoma phaeopygia*, commonly known as the Galapagos Petrel, is a long-lived procellariiform seabird endemic to the Galápagos Archipelago, where it breeds on the islands of Santa Cruz, San Cristóbal, Floreana, Santiago, and Isabela. The species is currently classified as Critically Endangered (Cruz-Delgado et al., 2010; BirdLife International, 2025).

The Galapagos Petrel was for a long time considered the same species as the Hawaiian Petrel (*Pterodoma sandwichensis*), being recognized as a unique species in the IUCN Red List in 2000 (Hilton-Taylor 2000) and officially accepted by the American Ornithologists ‘Union in 2002 (Banks et al, 2002). This followed various studies confirming significant differences in aspects such as morphology and vocalizations – Galapagos Petrels are generally larger, with longer wings and bills and distinct vocalizations – and significant genetic divergence, with no evidence found of gene flow between the Galapagos and Hawaiian groups (Tomkins and Milne, 1991; Browne et al., 1997).

The recognition of the Galapagos Petrel as a distinct species led to the elevation of its conservation status. Prior to the taxonomic split, *Pterodoma phaeopygia* was classified as Vulnerable in the 1994 IUCN Red List (Collar et al., 1994), with the Hawaiian population masking the severity of the situation for the Galapagos population. The Galapagos Petrel had already been exposed to long-standing threats, in some cases for centuries, from invasive species – such as black rats (*Rattus rattus*), cats (*Felis catus*), dogs (*Canis familiaris*), pigs (*Sus scrofa*) and fire ants (*Wasmannia auropunctata* and *Solenopsis geminata*) preying on eggs and chicks, as well as goats (*Capra hircus*), donkeys (*Equus asinus*) and cattle (*Bos taurus*) destroying nests and habitat – introduced during the distinct phases of human activity and colonization in the Galapagos (Cruz and Cruz, 1987; Causton et al., 2006; Phillips et al., 2012). Due to these pressures, the Galapagos Petrel population was already reported to be undergoing rapid decline and experiencing reproductive failure in the 1980s (Cruz and Cruz, 1987). The recognition of its status as a distinct species therefore highlighted its extremely vulnerable condition, leading to its classification as Critically Endangered in 2000 (Hilton-Taylor 2000).

Given the critical condition of the species, conservation efforts were already underway by 1982. These early interventions included the hunting of cats, pigs, goats, donkeys, and cattle during the petrel breeding season, the initiation of annual poison-based rodent control, and the establishment of the first large-scale monitoring efforts, which involved intensive searches for nests and the assessment of the petrels’ breeding success (Cruz and Cruz, 1987). Conservation initiatives have continued to expand over the decades. Since 2000, rangers, in collaboration with the Galápagos National Park, have conducted annual monitoring of petrel populations, nests, chicks and fledglings. They have also sustained efforts to control the black rat - arguably the most significant current threat to the species. Since 2020, the Galápagos Conservancy has partnered with field technicians and researchers to continue this work, ultimately yielding a robust 25-year dataset for Santa Cruz Island.

The present study constitutes the first comprehensive analysis of data collected over the past two and a half decades. It reveals long-term patterns in breeding, hatching, and fledging success, alongside the impacts of rodent control efforts on this Critically Endangered seabird population. This analysis is particularly timely; although introduced rats have been a recognized threat for decades, instances of rat predation on petrel nestlings continue to be documented as recently as 2025 (Tapia-Jaramillo et al., 2025). This highlights the persistence of severe anthropogenic pressures on the species and underscores the critical need for continued intervention.

## Methodology

### Study area

The monitoring was conducted in Santa Cruz Island, in the Galápagos Archipelago. In this area the Galapagos Petrel is known to restrict its nesting sites in high elevation areas largely within the borders of the protected National Park. These sites are typically located in the humid zone, generally above 200 meters, where deep soils allow for burrow excavation under native vegetation such as *Miconia robinsoniana* and ferns (Cruz and Cruz, 1987). Monitoring the Galapagos Petrel presents a serious logistical challenge due to the species’ pelagic lifestyle, the cryptic nature of its burrows, and the rugged, inaccessible vegetation surrounding them – often impenetrable thickets of native *Miconia* or the invasive blackberry (*Rubus niveus*). Furthermore, nests are frequently located on steep slopes, crater rims, or vertical ravine walls, making detection and access difficult and physically demanding (Cruz and Cruz, 1987; Cruz-Delgado et al., 2010). Consequently, monitoring key demographic parameters has proven challenging, and “new” nests – which were likely established previously – are frequently discovered when monitoring efforts are expanded.

### Field protocols

Due to these logistical challenges and successive years of new nest discovery, a revised protocol was established in 2016 whereby park rangers strictly monitored a unique number of marked nests. While doing so increased the efficiency of the monitoring process, it complicates the calculation of absolute population trends and necessitates analyses at the nest level. Consequently, the 25-year analysis in this report focuses on demographic indicators based on individual nest outcomes, specifically the number of chicks hatched and fledged per monitored nest. Following established protocols (Cruz and Cruz, 1987), successful fledging was inferred when a fully feathered chick reached the minimum expected fledging age and subsequently vacated the burrow with no associated signs of predation or in-nest mortality. Annual monitoring occurred consistently throughout the study period, with the exception of 2023, when no monitoring was conducted, and from 2020 to 2025 for which no fledgling data exists.

### Rodent Control Protocol

A rodent control protocol is applied annually to Santa Cruz Island to minimize the impact of invasive rodents on petrel breeding populations. The protocol consists of aerially distributing toxicant bait across the island, particularly during the dry season when rodents are more food deprived. The bait utilizes brodifacoum - a coumarin-based second-generation anticoagulant - which alters blood clotting ability and causes fatal hemorrhaging (Hoare and Hare 2006). Brodifacoum is the primary rodenticide used in island control protocols globally (Howard et al., 2007) and is preferred here due to its low bait shyness and high target susceptibility.

The year 2017 represented a distinct deviation from this standardized protocol. While black rats - a primary predator of petrels - were managed using this consistent methodology across all other years, 2017 was compromised by logistical disruptions in the bait supply chain. This led to delayed application and potential degradation of the active compounds, effectively leaving nests unprotected.

### Statistical analysis

Across the 25 breeding seasons on Santa Cruz Island (2000–2025), breeding productivity was assessed through repeated monitoring of a largely consistent sample of nest sites. To account for annual variations in monitoring effort, sampling effort is expressed as nest-years (one nest monitored for one breeding season). Annual effort ranged from 322 nest-years in 2000 to 760 nest-years in 2021, totaling 15,699 nest-years across the study period.

Because Galapagos petrels lay a single egg per clutch, three demographic variables were assessed as binomial proportions: (1) hatching success (the proportion of monitored nests that successfully hatched a chick), (2) breeding success (the proportion of monitored nests that successfully produced a fledgling), and (3) fledging success (the transition rate of hatched chicks that successfully fledged). The unit of analysis for statistical modeling was the annual breeding season, yielding a time series of yearly demographic estimates.

All statistical analyses were conducted in Python 3.10.4 (Python Software Foundation). Data aggregation was performed using pandas 1.5.3 (McKinney, 2010) and NumPy 1.24.3 (Harris et al., 2020). Generalized linear models (GLMs) were fitted utilizing statsmodels 0.14.0 (Seabold & Perktold, 2010), with patsy 0.5.3 (Smith et al., 2018) employed to generate basis matrices for the spline regressions. Graphical outputs were produced using Matplotlib 3.7.1 (Hunter, 2007).

To investigate temporal trends, GLMs were fitted using a binomial error distribution and a logit link function. Because the number of monitored nests varied annually, the binomial models were weighted by the number of trials (nests or chicks) per season, ensuring that years with greater sampling effort contributed proportionally more weight to the model estimations. To model non-linear interannual temporal fluctuations, we fitted time-varying models utilizing cubic B-splines with five degrees of freedom (df = 5). The significance of this temporal structure was evaluated by comparing the deviance of the spline models against a constant-mean (intercept-only) null model using likelihood ratio tests (*χ*^2^).

Colonial seabird demographic data frequently violate the assumption of independence, leading to extra-binomial variation. We formally assessed overdispersion by calculating the quasi-dispersion factor (*ϕ*), defined as the Pearson’s *χ*^2^ statistic divided by the residual degrees of freedom from the baseline models. Because sensitivity analysis revealed substantial overdispersion (*ϕ*> 1) - largely driven by the extreme 2017 management failure and inherent ecological clustering - we adopted a quasi-binomial approach for inference. Standard errors, confidence intervals, and p-values for all parameter estimates were conservatively inflated by scaling the variance-covariance matrix by the respective *ϕ*factor for each demographic rate.

## Results

### Hatching success

Across the 25-year study period, the effort-weighted mean hatching success was 0.657 chicks per nest-year, with a nearly identical unweighted annual mean (0.656). Excluding the 2017 rodent management failure year, the effort-weighted mean increased marginally to 0.669 (unweighted: 0.666). Because removing 2017 raised the long-term mean by a relative margin of only 1.8% (an absolute difference of 0.012 chicks per nest), demonstrating that this extreme low year does not disproportionately skew the long-term average, 2017 was retained in all subsequent models. Observed annual hatching success generally fluctuated near the long-term mean, peaking in 2005 (0.7159; 436 chicks from 609 nests) and dropping to its lowest recorded point in 2017 (0.426; 323 chicks from 759 nests). By the final available year (2025), hatching success had stabilized near the long-term average (0.650; 492 chicks from 757 nests).

Time-varying binomial models utilizing a cubic spline (df = 5) provided a substantially better fit than a constant-mean model (*χ*^2^ = 20.86; *p* < 0.001), confirming significant temporal variation. Sensitivity analysis revealed high overdispersion (*ϕ*= 12.88), largely driven by the 2017 management failure; excluding 2017 reduced the dispersion factor to *ϕ* = 4.12. Retaining 2017 and applying the full dispersion factor yielded conservative model-estimated trend values of 0.662 (95% CI: 0.495–0.797) in 2000, 0.682 (0.626–0.734) in 2010, 0.634 (0.581–0.684) in 2017, and 0.657 (0.547–0.752) in 2025. As expected, the actual observed value in 2017 dropped to 0.426 (323 chicks from 759 nests), falling substantially below the model-estimated mean for that year and corresponding directly with the absence of rodent control (Figure 1).

**Figure 1.**
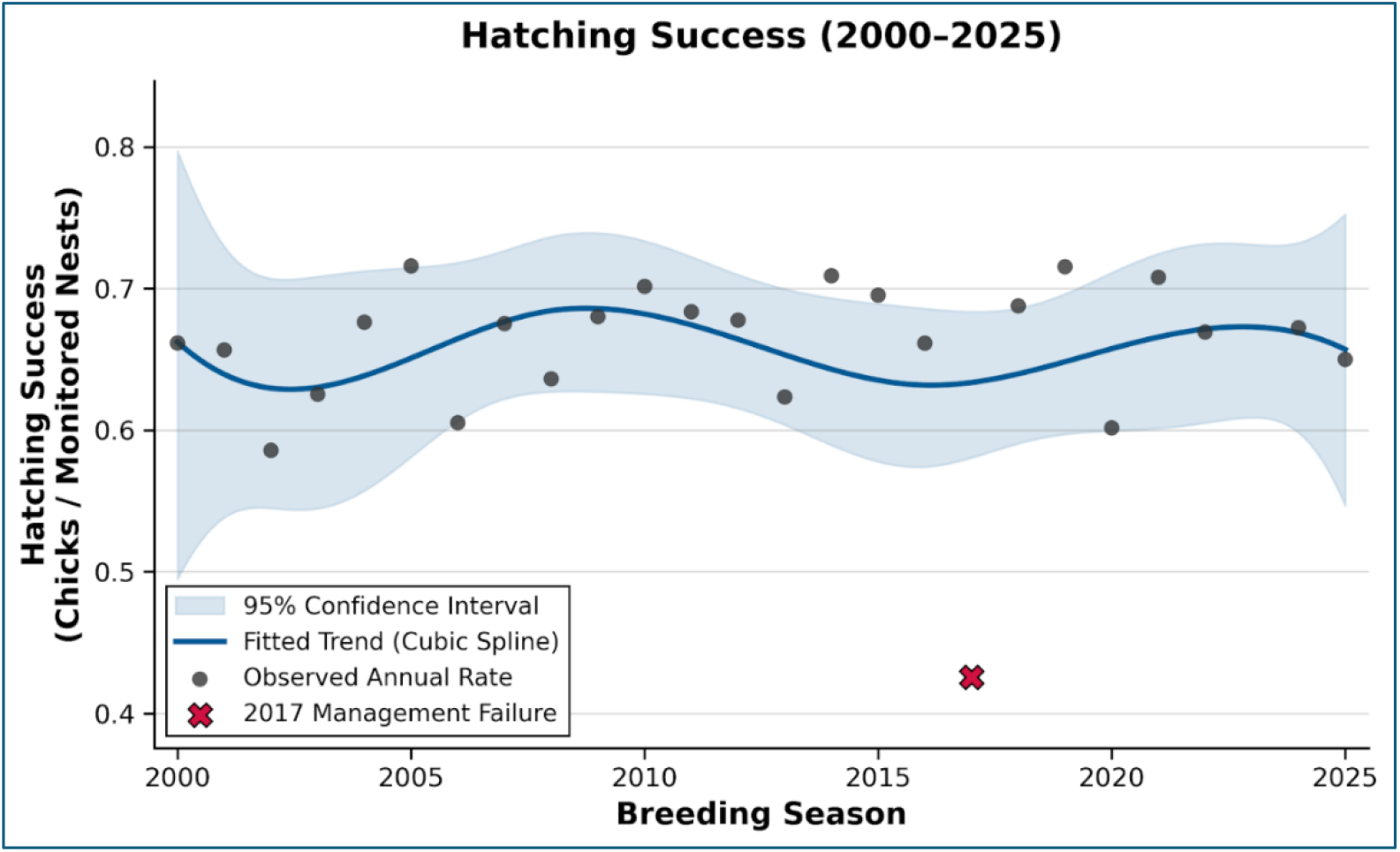
Galápagos petrel hatching success (chicks per monitored nest) from 2000 to 2025.

### Fledging Success

Fledging success - calculated as the transition rate of recorded chicks that subsequently left the nest - was generally high. The effort-weighted mean transition rate was 0.942, and the unweighted mean was also 0.942. The highest observed annual transition occurred in 2014 (0.989; 458 fledglings from 463 chicks), and the lowest occurred during the 2017 management failure year (0.864; 279 fledglings from 323 chicks), although the value remained relatively high.

Time-varying models (cubic spline, df = 5) fit substantially better than a constant-mean model (*χ*^2^ = 48.12, *p* < 0.001), indicating strong temporal structure in the chick-to-fledge transition (Figure 3). To account for year-to-year variability exceeding an ideal binomial process, uncertainty was conservatively inflated using a quasi-dispersion factor of *ϕ*= 10.13.

### Breeding Success

Data for breeding success (fledglings per monitored nest-year) were available for 20 breeding seasons (2000–2019). Across these seasons, monitoring effort summed to 11.908 nest-years, recording 7,352 chicks that successfully left the nest. The effort-weighted long-term mean breeding success was 0.617 fledglings per nest-year, with a similar unweighted mean of 0.617. Excluding the 2017 management failure year increased the effort-weighted mean to 0.634 and the unweighted mean to 0.631. Similar to hatching success, this exclusion shifted the long-term estimate upward by a relative margin of only ∼2.8% (an absolute difference of 0.017 fledglings per nest). Because this extreme year does not fundamentally alter the long-term baseline, 2017 was retained in all subsequent breeding success models.

Observed breeding success peaked in 2014 (0.701; 458 fledglings from 653 nests) and was markedly lower than all other years in 2017 (0.368; 279 fledglings from 759 nests). In the final year available for this metric (2019), breeding success remained high (0.688; 522 fledglings from 759 nests), following a preceding year (2018) that was close to the long-term mean (0.605; 459 fledglings from 759 nests).

Time-varying models (cubic spline, df = 5) fit substantially better than a constant-mean model (*χ*^2^ = 80.94, *p* < 0.001), indicating strong temporal structure (Figure 2). Similar to hatching success, year-to-year variability exceeded that expected under an ideal binomial process (*Φ* = 13.65). Sensitivity analysis confirmed this overdispersion was largely driven by 2017; excluding it reduced the dispersion factor to *ϕ* = 3.96. Retaining 2017 and utilizing the full dispersion factor resulted in model-estimated mean breeding successes of 0.615 (95% CI: 0.432–0.770) in 2000, 0.666 (0.609–0.719) in 2010, 0.544 (0.469– 0.617) in 2017, and 0.660 (0.540–0.762) in 2019. The 2017 observed breeding success rate exhibited the greatest negative divergence from the model-estimated trend.

**Figure 2.**
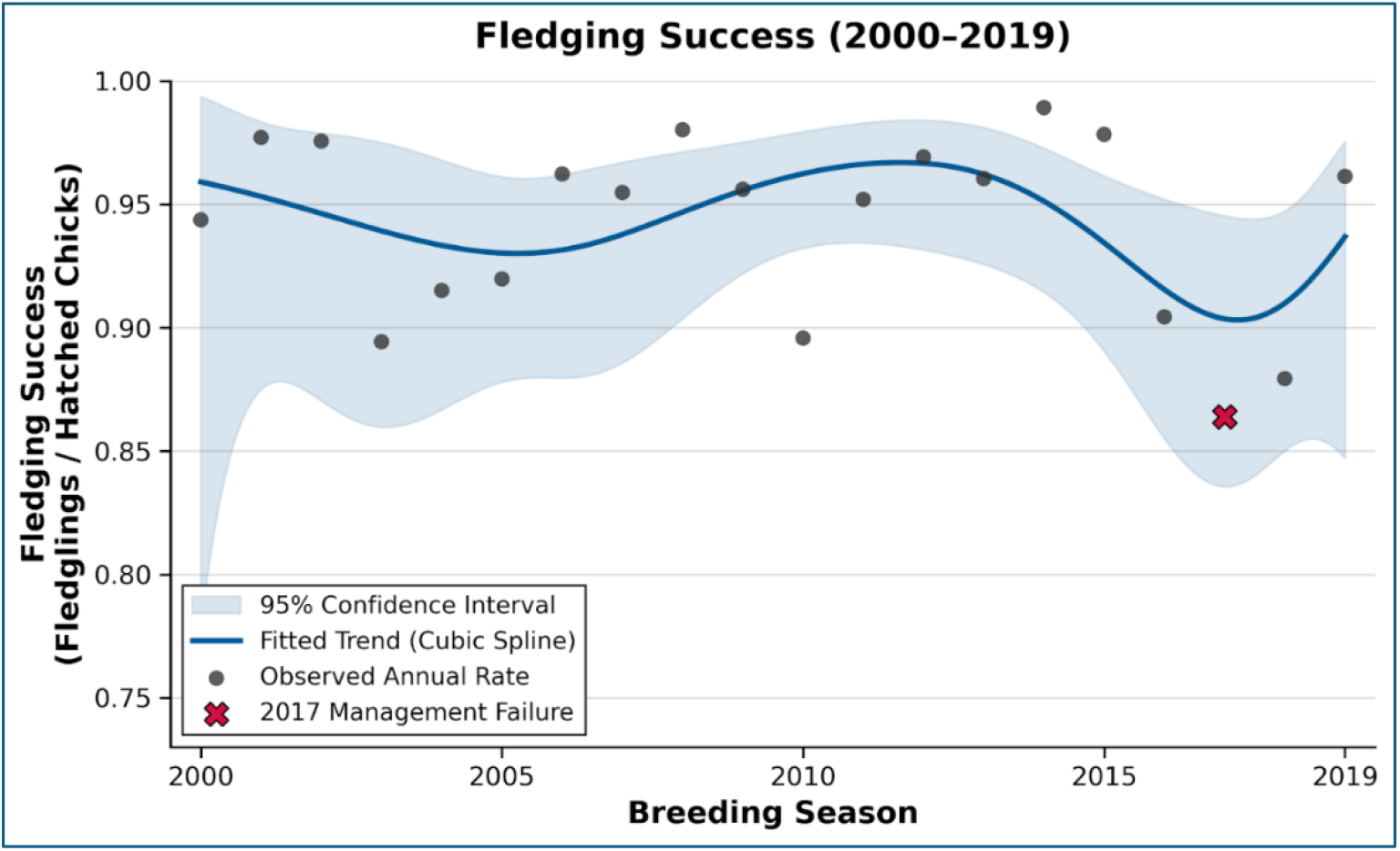
Galápagos petrel fledging success (fledglings per hatched chicks) from 2000 to 2019.

**Figure 3.**
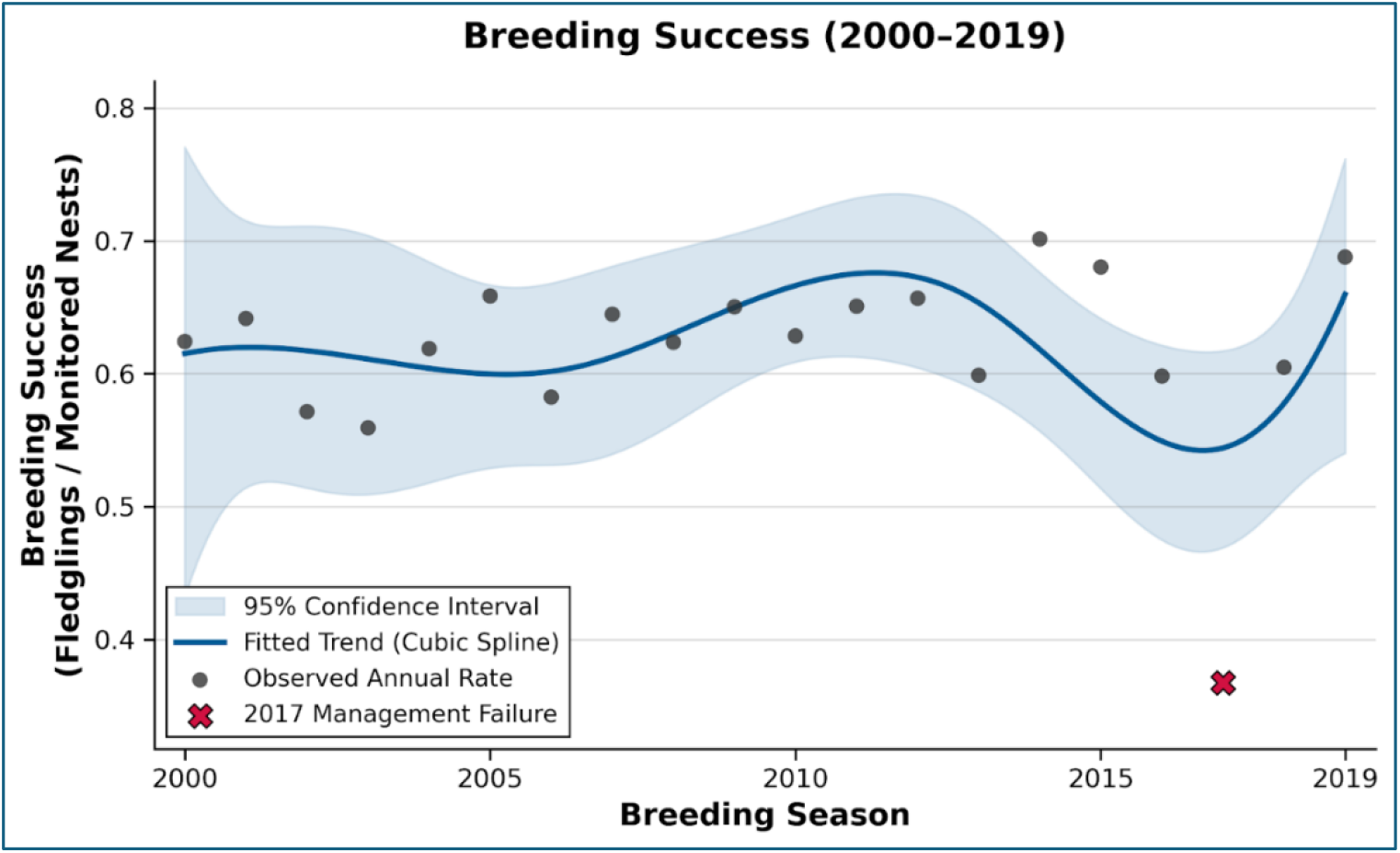
Galápagos petrel overall breeding success (Fledglings per monitored nests) from 2000 to 2019.

## Discussion

The Galapagos Petrel exhibited little variation in hatching, breeding and fledging success over 25 years (2000-2025), showing positive resilience against strong climatic events such as El Niño, which frequently disrupt biodiversity across the Galápagos Archipelago (Vargas et al., 2006; Dubiner et al., 2025). On the other hand, a single year of lapsed rodent control in 2017 immediately severely affected the seabirds hatching and breeding success, emphasizing the persistent impacts of invasive rodent species and the need for continued control. Fledging success does not seem to be significantly affected by the year without rodent control, indicating that the threat of invasive rodents is possibly significantly higher during egg incubation and early chick-rearing phases rather than when chicks are already older and nearer fledging stage.

Procellariiform seabirds exhibit extreme life-history traits associated with *K*-selection, characterized by delayed sexual maturity, exceptionally high annual adult survival, and profound parental investment in a single-egg clutch per breeding attempt (Hamer et al., 2001; Warham, 1990). Because of their longevity, population stability in these taxa does not require near-perfect annual reproductive output. Warham (1990) established that an overall breeding success of approximately 50% is generally sufficient to maintain a stable petrel population, whereas optimal, predator-free environments frequently produce success rates of 70% or greater. Over the 25-year period analyzed in this study, the Galápagos Petrel (*Pterodroma phaeopygia*) exhibited a hatching success of 66%, a fledging success of 94%, and an overall breeding success of 62%. These metrics sit comfortably within the established demographic thresholds for a healthy and stable population; notably, one year even reached the optimal threshold of 70% breeding success (2014) and other two years also approached this high mark (2015, 2019). Comparing these demographic parameters to other procellariiform populations globally also indicates that our recorded values fall well within the typical range for the taxon. For instance, a study on the Tahiti Petrel (*Pseudobulweria rostrata*) in New Caledonia, conducted in an area free from invasive species threats, reported 61% hatching, 81% fledging, and 50% overall breeding success (Pagenaud et al., 2024), despite a limited sample size of assessed burrows. Notably, these values from a predator-free environment are lower than those of the Galapagos petrel - a species threatened by the invasive black rat for the last decades - thereby highlighting the high efficacy of the rodent management programs currently implemented across the archipelago.

Similar reproductive trends are observed in other procellariiform populations experiencing minimal to no invasive rat predation. For example, Leach’s Storm-Petrels (*Hydrobates leucorhous*) on Great Island, Newfoundland, showed 72% hatching and 71% overall breeding success in 2000 (Stenhouse & Montevecchi, 2000), and more recently, in Mexico, demonstrated 80.5% hatching, 87% fledging, and 70% overall success between 2014 and 2015 (Bedolla-Guzmán et al., 2017). Other comparable metrics include the Bermuda Petrel (*Pterodroma cahow*), with 68% hatching, 91% fledging, and 62% breeding success (Madeiros et al., 2012); Cory’s Shearwater (*Calonectris borealis*) in the Azores, Portugal, at 51%, 87%, and 45% for the same respective metrics (Ramos et al., 2003); and Scopoli’s Shearwater (*Calonectris diomedea*) in a low-threat area in Greece, which presented approximately 77% hatching, 87% fledging, and 67% overall breeding success (Karris et al., 2024).

Collectively, these unthreatened or low-threat procellariiform populations yield mean reproductive success rates of approximately 68% for hatching, 87% for fledging, and 61% for overall breeding. Against these baselines, the Galapagos petrel sits merely 2% below the hatching mean, yet 7% above the fledging mean, resulting in a marginal 1% difference in overall breeding success. This is maintained even when accounting for the catastrophic 2017 rodent management failure. This underscores that over the past 25 years, the Galapagos petrel has exhibited robust demographic resilience and positive reproductive outputs comparable to numerous unthreatened procellariiform populations.

However, these values must be interpreted with methodological caution. Because our monitoring protocol consecutively assessed the same nests over time, there is a possibility that the success rates are slightly inflated. Procellariiforms typically exhibit high levels of nest-site tenacity and mate fidelity, behaviors that enhance parental coordination and subsequently improve breeding success (Warham, 1990). Conversely, breeding failure frequently triggers divorce as individuals seek better breeding conditions (Thibault, 1994; Bourgeois et al., 2014). For instance, in the Yelkouan Shearwater (*Puffinus yelkouan*), faithful breeders achieved a 67.5% breeding success rate within a given year, whereas movers attained only 43.8%, alongside a 97.3% nest-cavity fidelity rate following a successful breeding attempt (Bourgeois et al., 2014). Given these behavioral dynamics, it is reasonable to hypothesize that the approximately 760 nests monitored regularly in our study area are disproportionately occupied by experienced, highly successful adults that have established optimal mate and nest pairings. This is further supported by a mean nest occupancy rate of 93% over the last five years, suggesting a well-established subpopulation on Santa Cruz Island. Consequently, these metrics may not fully represent the reproductive reality of other unmonitored or newly established nesting areas, highlighting the need for broader spatial monitoring. Nonetheless, the overall reproductive metrics remain highly optimistic for the conservation of the endangered Galapagos petrel.

Over the quarter-century study period, the only anomalous collapse in Galapagos petrel reproductive success coincided directly with the 2017 rodent management failure. During this single season, hatching success plummeted to 43% and overall breeding success to 37%, representing relative decreases of approximately 35% and 40% from the long-term baseline, respectively. The overall breeding success during this year (37%) is similar to that of other Procellariids affected by the presence of invasive rats (32 – 36%) (Pagenaud et al., 2024; Bester et al., 2007). Fledging success was less severely impacted, maintaining a comparatively high 86% rate; however, this still constituted the lowest recorded value for this metric and an 8.5% drop from the long-term mean. This demographic contrast underscores the critical necessity of uninterrupted predator control in insular ecosystems.

The severe impact of invasive mammalian predators on procellariiforms is well documented globally. For example, on Marion Island, Great-winged Petrels (*Pterodroma macroptera*) threatened by feral cats exhibited breeding success rates ranging from 0% to 20.5% between 1979 and 1984. Following successful predator control, breeding success rebounded and stabilized at approximately 60% (Cooper & Fourie, 1991) - a baseline similar to the normal mean discussed previously. However, invasive rodents present a particularly complex management challenge, affecting an estimated 80% of the world’s archipelagos (Atkinson, 1985). Complete eradication of rodents is notoriously difficult, particularly on tropical islands (both wet and dry) compared to their temperate counterparts (Keitt et al., 2014). An analysis of 216 rodenticide-based operations revealed that eradications are significantly more likely to fail on islands with a mean annual temperature of 24°C or higher and unpredictable rainfall patterns (Holmes et al., 2015). Santa Cruz Island matches both criteria precisely, with an annual mean air temperature of 24.14°C and a 60% Coefficient of Variation in mean precipitation between 2000 and 2025. In warmer climates, higher natural food availability can reduce bait consumption, while unpredictable precipitation obscures the optimal timing for control measures by masking nonseasonal peaks in rodent productivity (Holmes et al., 2015). Although non-target native species can sometimes compete for bait and reduce program effectiveness (Griffiths et al., 2011), this specific interference appears absent in the Galapagos (Holmes et al., 2015). Even without bait competition, the remaining factors continue to make rodent extirpation in tropical systems challenging. A single surviving pregnant female might be able to rapidly repopulate an island (Witmer et al., 2011; Valdez et al., 2018). Furthermore, multi-paternity in *Rattus rattus* populations ensures that even small founder groups maintain significant genetic variation, allowing recovering populations to quickly bounce back from incomplete control attempts (King et al., 2013).

While achieving full extirpation on islands with these climatic characteristics is exceptionally challenging, our data suggest that sustained annual rodent control - even without achieving total eradication - is sufficient to maintain petrel reproductive success at healthy thresholds, albeit requiring continued investment. This aligns with findings from the Chafarinas Islands, which hosted high densities of invasive black rats until an intensive anticoagulant control campaign was initiated between 1999 and 2004. Following control, Cory’s Shearwater breeding success surged from 44% in 1997 to 70% in 2000, stabilizing at 71% through 2004, effectively mirroring rat-free colonies (Igual et al., 2005).

A parallel dynamic happens to the Galapagos petrel. Although black rats remained a persistent background threat throughout our 25-year study period, active management sustained overall breeding success at a robust 63%. The unintentional 2017 management failure effectively served as a natural quasi-experiment, confirming that the absence of rodent control immediately triggers a severe demographic decline, exactly as seen in previous case-studies. Ultimately, these findings solidify the hypothesis that, even when permanent eradication in tropical ecosystems remains difficult, sustained annual rodenticide programs are highly effective and able to keep vulnerable petrel populations at an healthy and even close to optimal reproductive success threshold.

The observation that fledging success was significantly less affected by the catastrophic 2017 management failure also provides valuable insights into the ecological dynamics of invasive rodent predation. Multiple studies demonstrate that procellariiform populations are acutely vulnerable to predation during the egg phase and early post-hatching period. For instance, chicks are most frequently preyed upon between 5 and 7 days of age (Thibault, 1995) or between 2 and 7 days (Igual et al., 2006), while temporary egg neglect by parents can rapidly result in substantial rat predation (Imber, 1984). However, once chicks cross a developmental threshold - such as reaching three weeks of age or two-thirds of adult weight, as observed in Cory’s Shearwater - predation risk decreases significantly or ceases entirely (Igual et al., 2006; Thibault, 1995).

Our data strongly indicate that this stage-specific vulnerability applies directly to the Galapagos petrel. Fledging success remained notably high during the 2017 management lapse, even as hatching success collapsed due to the unchecked rodent population. Consequently, it is reasonable to hypothesize that the Galapagos petrel experiences a severe predation bottleneck during egg incubation and the early chick-rearing phase. Because the phenology of these reproductive stages is already well-documented for the species (Harris, 1970; Cruz & Cruz, 1990), conservation efforts can be strategically targeted to ensure that rodent populations are most aggressively suppressed during these specific temporal windows. Our results indicate stage-specific vulnerability in the egg and early chick-rearing phases, with older chicks showing greater resilience, providing a basis for optimizing rodent management strategies also for other threatened procellariiforms.

## Future directions and limitations

The present study acknowledges certain methodological limitations inherent to the long-term monitoring of petrel populations. As previously noted, the monitored nests were not randomly selected, and the same subset of burrows was assessed consistently across the 25-year dataset. This approach introduces potential biases regarding population representativeness and spatial extrapolation (Hunter, 2001; Baker et al., 2010). To mitigate these issues, future research on the Galapagos petrel should aim to incorporate randomized sampling designs - such as stratified transects across newly identified subcolonies (Fewster et al., 2009) - to capture broader demographic trends. While we recognize that the feasibility of such randomization remains highly dependent on the accessibility and spatial clustering of burrows across the rugged island terrain, strictly targeted sampling should ideally be phased out where logistically possible (Parker and Rexer-Huber, 2020).

Furthermore, expanding the scope of environmental and behavioral data collected during nest visits will enable a more granular analysis of the variables driving reproductive success. Studies on other procellariiforms that incorporate microhabitat preferences, nesting typologies, nest-site and mate fidelity, sociality, specific depredation patterns, and diet (Thibault, 1994; Stenhouse and Montevecchi, 2000; Bedolla-Guzmán et al., 2017; Karris et al., 2024) have yielded critical insights for species conservation.

For the Galapagos petrel, detailed monitoring of the entire reproductive cycle - from egg morphometrics to distinct chick growth stages (Ramos et al., 2003) - is particularly crucial. Because our findings strongly suggest a specific size or developmental threshold at which chicks escape peak predation risk, pinpointing this precise milestone could refine and optimize targeted conservation interventions for this endangered seabird.

Lastly, Galapagos petrels exhibit notable plasticity in their nest-site selection, utilizing microhabitats that range from relatively open terrain to deep vegetative cover. As demonstrated in other procellariiform populations, such microhabitat variations can act as a significant determinant of overall breeding success (Stenhouse & Montevecchi, 2000). Furthermore, island-based conservation trials have shown that the implementation of artificial burrows can result in higher reproductive success rates than natural nests (Oliveira et al., 2020). Given that nest quality and subsequent site fidelity are critical drivers of procellariiform reproductive output (Bourgeois et al., 2014), experimental deployments of artificial burrows warrant consideration. Testing the efficacy of these structures - particularly in subcolonies disproportionately affected by breeding habitat degradation, human disturbance, or interspecific competition - represents a promising, proactive conservation strategy to safeguard an endangered species such as the Galapagos petrel.

Despite the potential for habitat availability bias and extrapolation limitations, the extensive timeframe and substantial sample size of this targeted dataset provide robust, highly optimistic insights into the conservation trajectory of the Galapagos petrel. Our findings align closely with the broader procellariiform literature, confirming that while tropical island ecosystems present unique and formidable challenges for invasive rodent eradication, sustained management can successfully secure viable reproductive outputs. Ultimately, this study significantly advances the evidence base for island petrel conservation, offering actionable perspectives for mitigating rodent impacts in complex tropical insular environments.

## Conclusion

Our results demonstrate that, although the Galapagos petrel remains endangered, its reproductive output can remain remarkably stable over decadal timescales under sustained management. However, this demographic stability is highly dependent on continuous rodent control, as even a single year of management failure can cause catastrophic declines in breeding success. These findings highlight the disproportionate vulnerability of early life stages to invasive predators, particularly during the incubation and initial chick-rearing phases. More broadly, our study underscores the critical importance of uninterrupted control efforts on tropical islands, where complete rodent extirpation is often difficult. Extending these management strategies across the Galápagos archipelago - particularly to less-studied populations on neighboring islands - will be essential to ensure the species’ long-term persistence. Ultimately, this 25-year dataset serves as a powerful model for successful procellariiform conservation in similarly invaded island ecosystems worldwide.

